# A conservation assessment of the freshwater gastropods of South Dakota based on historical records and recent observations

**DOI:** 10.1101/309385

**Authors:** Bruce J. Stephen

**Author notes:** Current Address: Acadia Institute of Oceanography, Seal Harbor, ME 04675. Correspondence: Bruce J Stephen.

## Abstract

In South Dakota, like most U.S. states, up-to-date knowledge of the distribution of freshwater gastropod species is lacking and historical data suffers from a host of synonyms. I consulted literature records and online museum databases to compile a list of freshwater gastropods historically recorded for South Dakota. I used systematic studies and regional records to evaluate each historically-listed species reducing 54 nominal species to 25 expected to inhabit South Dakota. This, along with recent survey data from across the state, enable a benchmark conservation status to be established for the freshwater gastropods of South Dakota. My preliminary conservation evaluation indicates *Planorbula armigera* is critically imperiled (S1), while three species; *Ferrissia rivularis, Campeloma decisum*, and *Amnicola limosus* are imperiled (S2). The status of historical species not observed recently, and suspected inhabitants known from adjacent states are discussed.

## Introduction

Freshwater snails are important components of aquatic systems. Snails consume periphyton and play a role in the breakdown of leaf litter (Dillon 2000, Brady and Turner 2010). Snails are eaten by a variety of invertebrate and vertebrate species, particularly fishes and waterfowl (Osenberg 1989, Brönmark and Vermaat 1998, Swanson and Duebbert 1989, Dillon 2000, Tiemann et al. 2011). Competitive and more complex interactions are found between freshwater snails and tadpoles (Brönmark et al. 1991, Smith et al. 2012). Complex interactions also exist among snails, fish, and epiphyton (Brönmark and Vermaat 1998). A large amount of energy throughput in aquatic systems is via snails (Newbold et al. 1983, Richardson et al. 1988, Brown 2001). Snails host the early life stages of a variety of trematode parasites (Dillon 2000). Despite being important ecosystem components, information about the distribution and abundance of aquatic snails is lacking.

In South Dakota, specific information on the distribution of aquatic gastropods comes primarily from just a few studies (Over 1915a, Over 1915b, Over 1928, Henderson 1927). Recent work on South Dakota freshwater gastropods is focused on few or non-indigenous species (Hershler 1996, Anderson 2004). South Dakota is not an anomaly; current information on freshwater snail distribution across the U.S. is missing for many states. Most of the information about freshwater snail distribution in North America comes from works that are broad in scope covering the whole of North America (Burch 1989, Johnson *et al*. 2013). There is a need to compile region-specific data in order to evaluate the conservation status of freshwater snails.

The taxonomy of freshwater snails is in flux and authors use a variety of names for the same species. I follow the taxonomy in Thorp and Rogers (2016).

A list of fresh water gastropod species is produced herein based on the historical literature and museum collections updated with taxonomic revisions. Regional studies from the adjacent states of Iowa, Nebraska, North Dakota, and Wyoming are used to suggest regional species abundance. However, as in South Dakota, current surveys are almost nonexistent. The historical records act as a base to compare data from 98 recent survey sites that will inform changes in species distribution and provides preliminary conservation status to the freshwater snails of South Dakota.

## Methods

Species presence records were obtained from published literature and online museum collections. There are few studies that focus on South Dakota freshwater snails. Most information comes from just a few sources (Over 1915a, Over 1915b, Over 1928, Henderson 1927).

Online museum catalogs for The Florida Museum of Natural History (FLMNH), Illinois Natural History Survey Mollusk Collection (INHS), The Academy of Natural Sciences of Drexel University (formally The Academy of Natural Sciences Philadelphia) (ANS), and Harvard Museum of Comparative Zoology (MCZ) were searched for records of freshwater snails from South Dakota. The Academy of Natural Sciences of Drexel University (ANS), and Harvard Museum of Comparative Zoology (MCZ) had collection from South Dakota. Appendix A is the database created from these sources.

Historical species listed in literature and museum records were scrutinized to assess the likelihood of identification errors based on the presence of species in other states within the region. The primary works examined from each state were North Dakota (Cvancara 1983), Nebraska (Stephen 2015), Wyoming (Beetle 1989), and Iowa (Stewart 2006). Updating and merging of species names were done by following taxonomic specific works of families as follows: Planorbidae (Hubendick and Rees 1955); Physidae (Dillon et al. 2002, Dillon and Wethington 2004, Wethington and Lydeard 2007); Lymnaeidae (Hubendick 1951, Remigio 2002, Correa et al. 2010); Ancylidae (Walther et al. 2006, Walther et al. 2010). Where no recent studies have been completed I follow the comprehensive guide by Burch (1989). A brief narrative description, including synonyms used in South Dakota studies, is produced for every species historically or currently present in South Dakota.

Recent data comes from surveys conducted throughout South Dakota between 2005 and 2011. Sample sites were chosen systematically while driving through the region. I focused on areas of high wetland density or wetland complexes such as sections of the Prairie Potholes and Middle Rockies ecoregions but also covered the majority of the state sampling within each major level III EPA ecoregion (Bryce et al 1998) and 46 of 66 counties. Some sample sites were chosen from historical literature; this included haphazard sampling in regions without specific historical locality data.

Sampling was done using dip nets, hand nets, and by hand, and included visual examination of shorelines, bottom substrate, vegetation, detritus, and shallow water structures. Searching continued until 30 minutes had elapsed since finding an additional species.

I indicate historical distribution for each species that had specific county records by shading the county on maps (Figures 2–5). Recent survey observations are included in Figures 2–5 as open circles at the approximate sample site.

**Figure 1.**
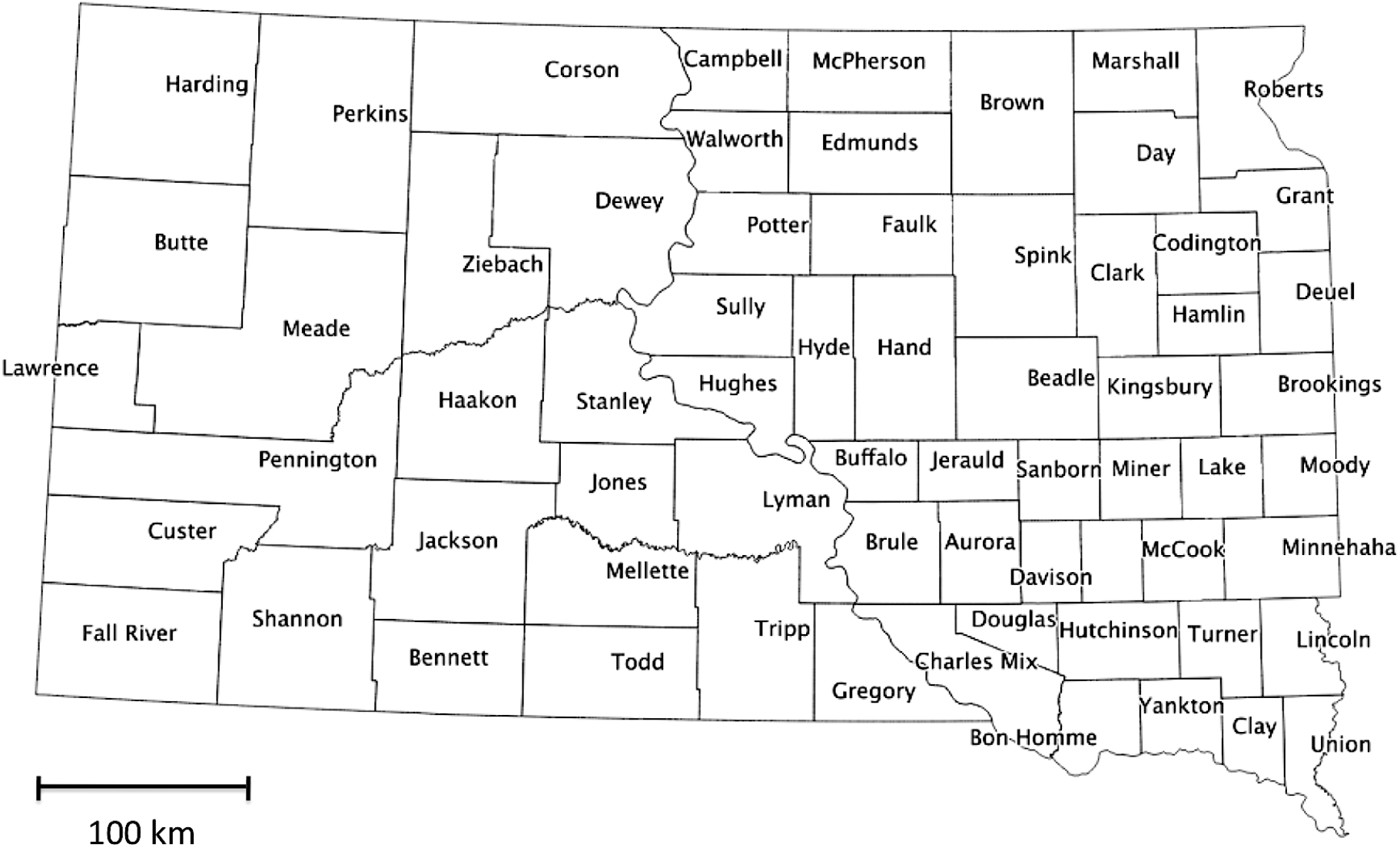
Map of South Dakota’s 66 counties.

**Figure 2.**
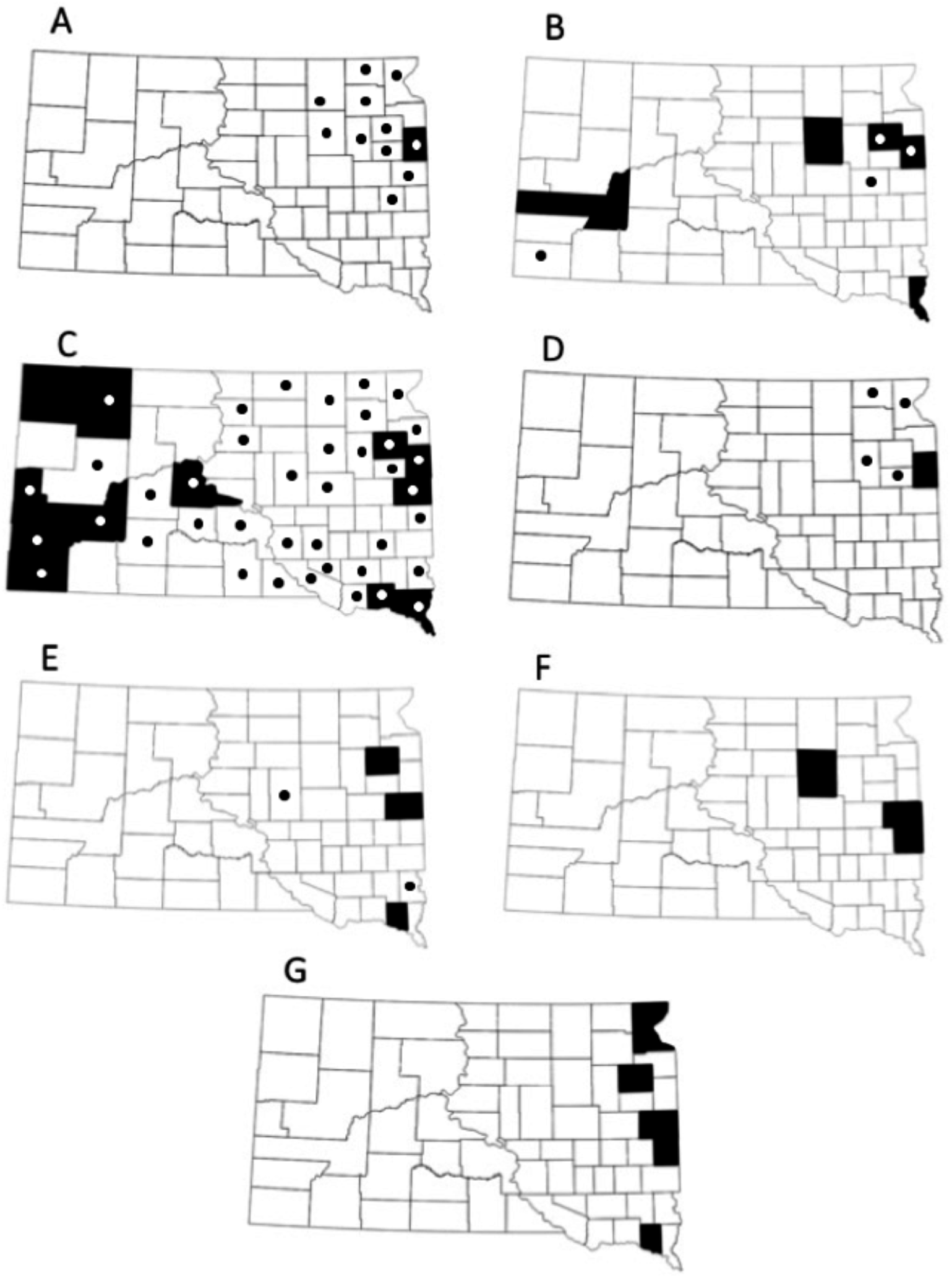
County level locality for (A) *Aplexa elongata*, (B) *Physa acuta*, (C) *Physa gyrina*, (D) *Physa jennessi*, (E) *Amnicola limosus*, (F) *Cincinnatia integra* and (G) *Probythinella emarginata*. Dark shading indicates counties with historical records, circles indicate counties with observations made during the current survey.

**Figure 3.**
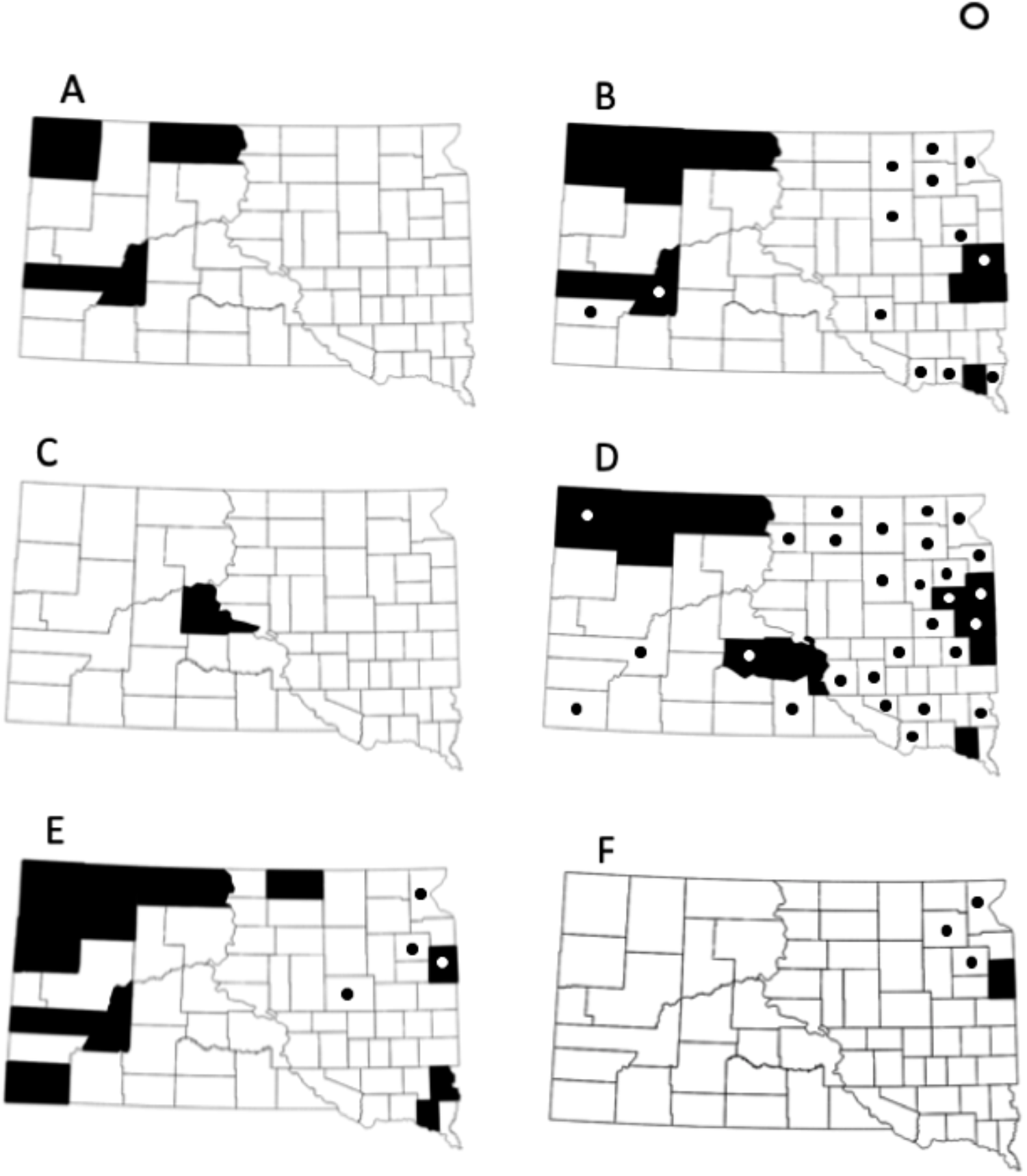
County level locality for (A) *Lymnaea bulimoides*, (B) *Lymnaea caperata*, (C) *Lymnaea catascopium*, (D) *Lymnaea elodes*, (E) *Lymnaea humilis*, and (F) *Lymnaea stagnalis*. Dark shading indicates counties with historical records, circles indicate counties with observations made during the current survey.

**Figure 4.**
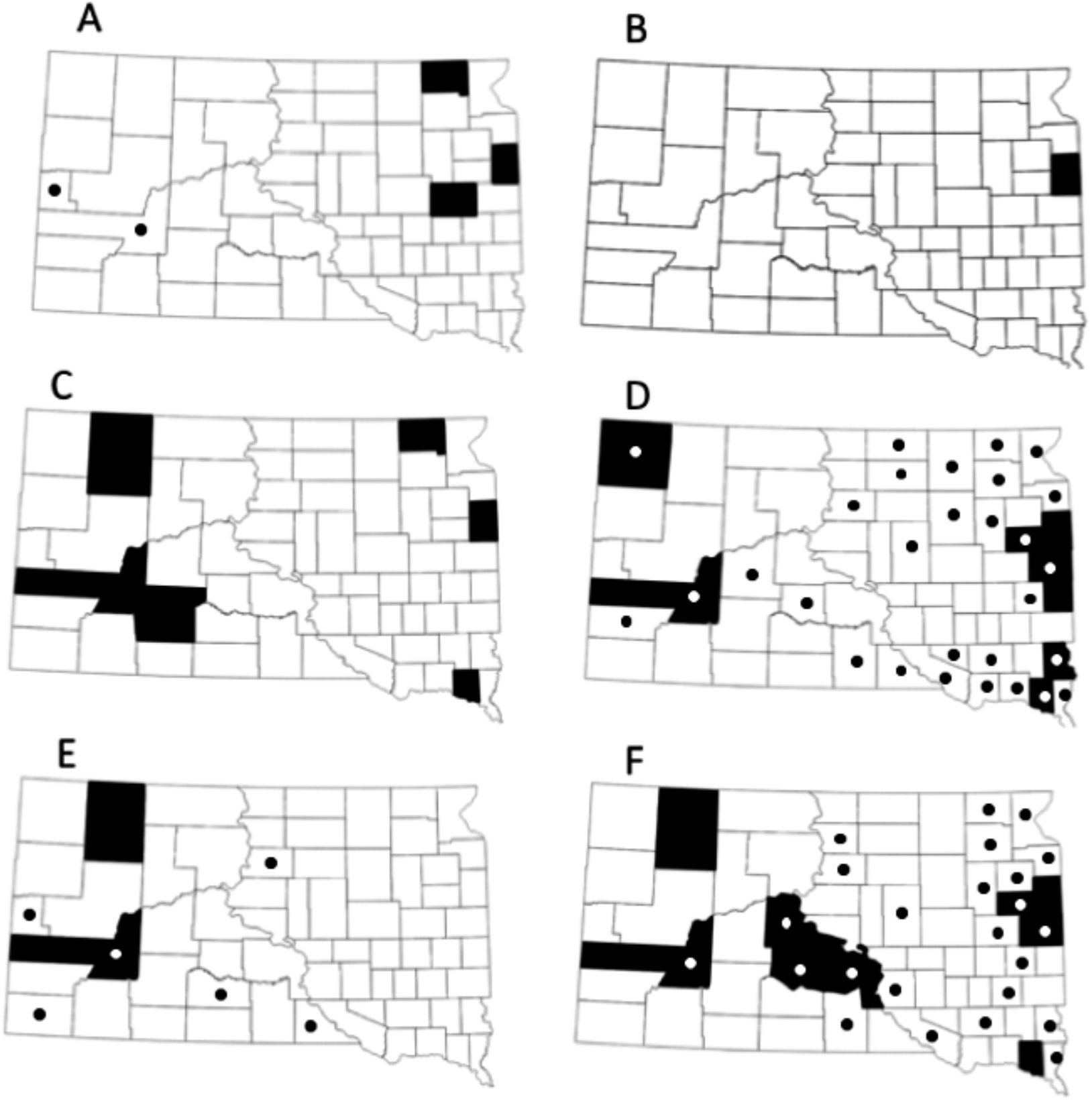
County level locality for (A) *Ferrissia rivularis*, (B) *Gyraulus crista*, (C) *Gyraulus deflectus*, (D) *Gyraulus parvus*, (E) *Helisoma anceps*, and (F), *Helisoma trivolvis*. Dark shading indicates counties with historical records, circles indicate counties with observations made during the current survey.

**Figure 5.**
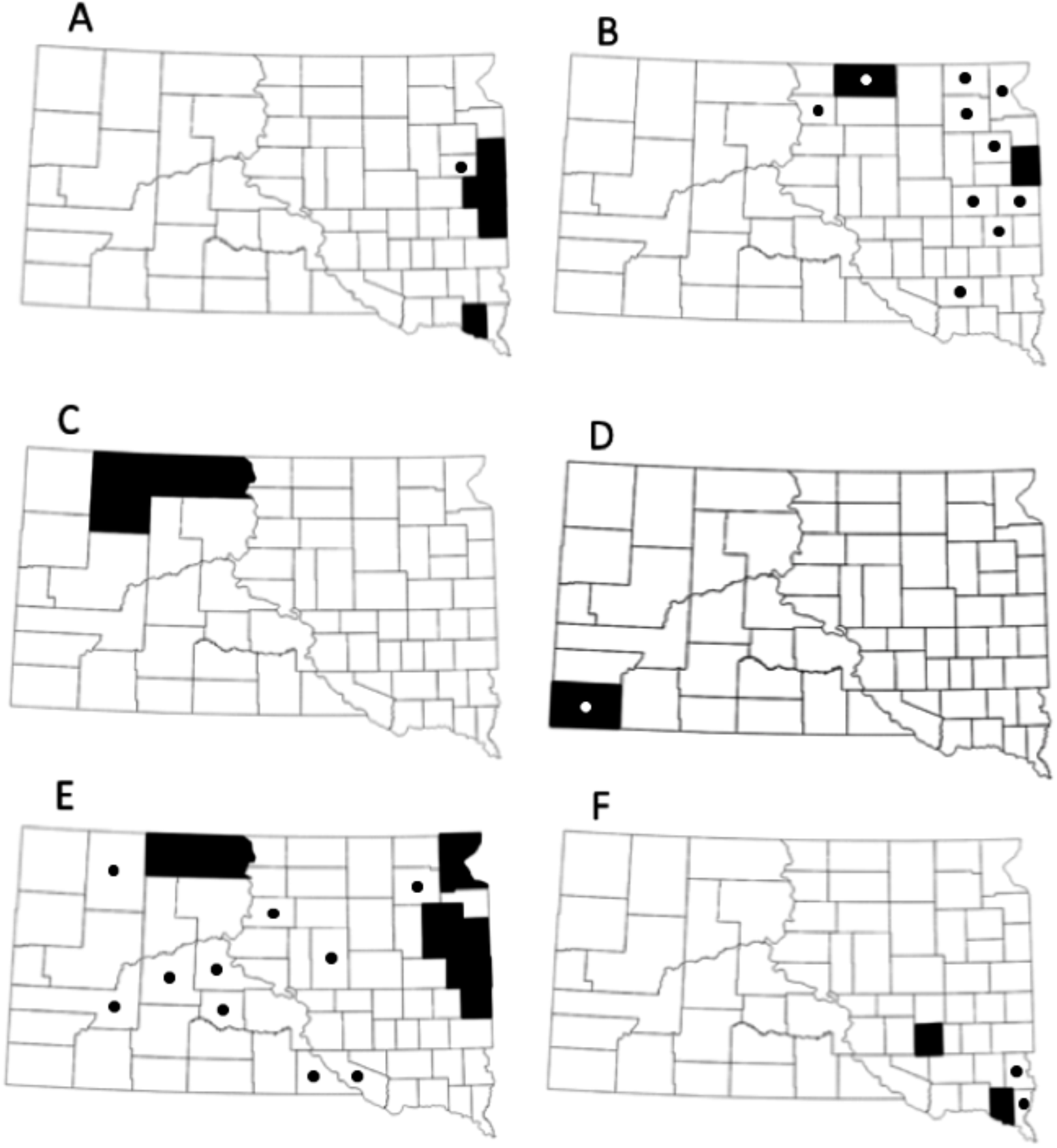
County level locality for (A) *Planorbula armigera*, (B) *Promenetus exacuous*, (C) *Promenetus umbilicatellus*, (D) *Melanoides tuberculatus*, (E) *Valvata tricarinata*, and (F) *Campeloma decisum*. Dark shading indicates counties with historical records, circles indicate counties with observations made during the current survey.

Conservation status was determined using species incidence of recent survey data via a modified quartile method (Gaston 1994, Dillon 2013) to separate species into local conservation ranks as developed by the Nature Conservancy and NatureServe (Master *et al*. 2009). Following this, the 5% of species with the lowest incidence are designated S1, critically imperiled locally. The 20% left in this quartile are S2, imperiled locally. The second quartile are S3, uncommon, the third S4, apparently secure, and the fourth quartile are S5, demonstrably secure. Some adjusting of quartiles was done to keep species with the same incidence value within the same conservation status. This resulted in a different number of species in each quartile.

Species from historical studies and museum records not observed recently are categorized as either SU - Unable to evaluate. (Taxonomically suspect or historical identification error) or SH - Possibly extirpated (historical listing only). Herein I also discuss other species that may inhabit South Dakota but did not rank them unless specific records exist of their presence within the state.

## Results

My literature and online review of the freshwater snails of South Dakota returned 214 records and names of 54 freshwater snail species. I reduced these 54 names to 25 valid species primarily by grouping synonyms (Table 1). My recent survey data from 98 water bodies yielded 18 species and 301 records. Species are ranked by incidence and given conservation status (Table 2) and county level distributions are mapped (Figs 2–5); shaded counties indicate literature records when a specific county could be determined. The approximate locale of recent species observation is indicated with open circles. Samples sites contained from 0-6 species with an average of 3.1 species per site. Snails were not observed at three sample sites, two reservoirs within the Black Hills and one reservoir in the extreme northwest within the Northwestern Great Plains ecoregion. Both reservoirs within the Black Hills (Angostura Reservoir and Roubaix Lake) had recently gone through restoration projects and were being filled after recent water level reductions. The third reservoir (Lake Gardner) had a pH 10.

**Table 1.**
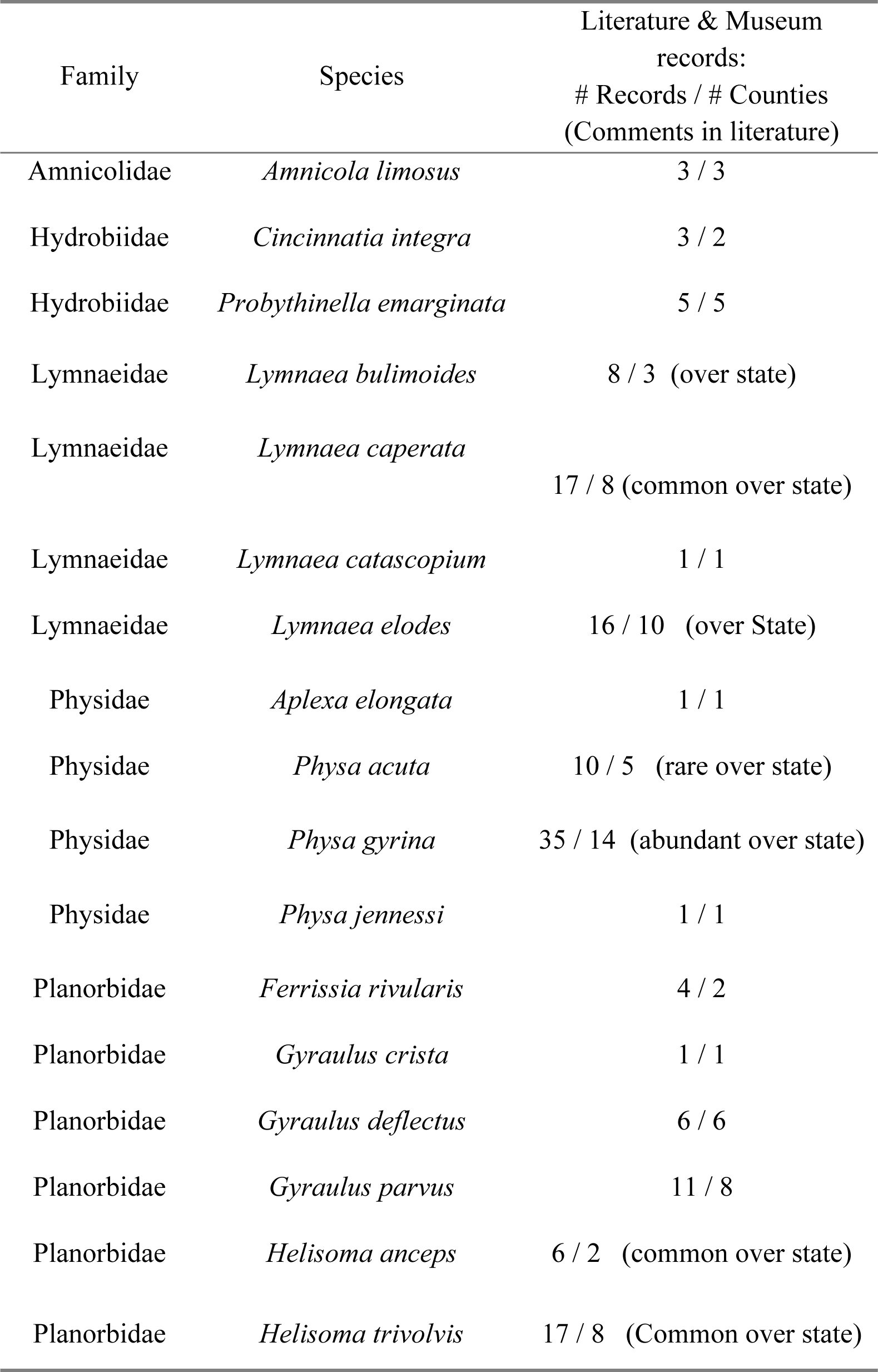
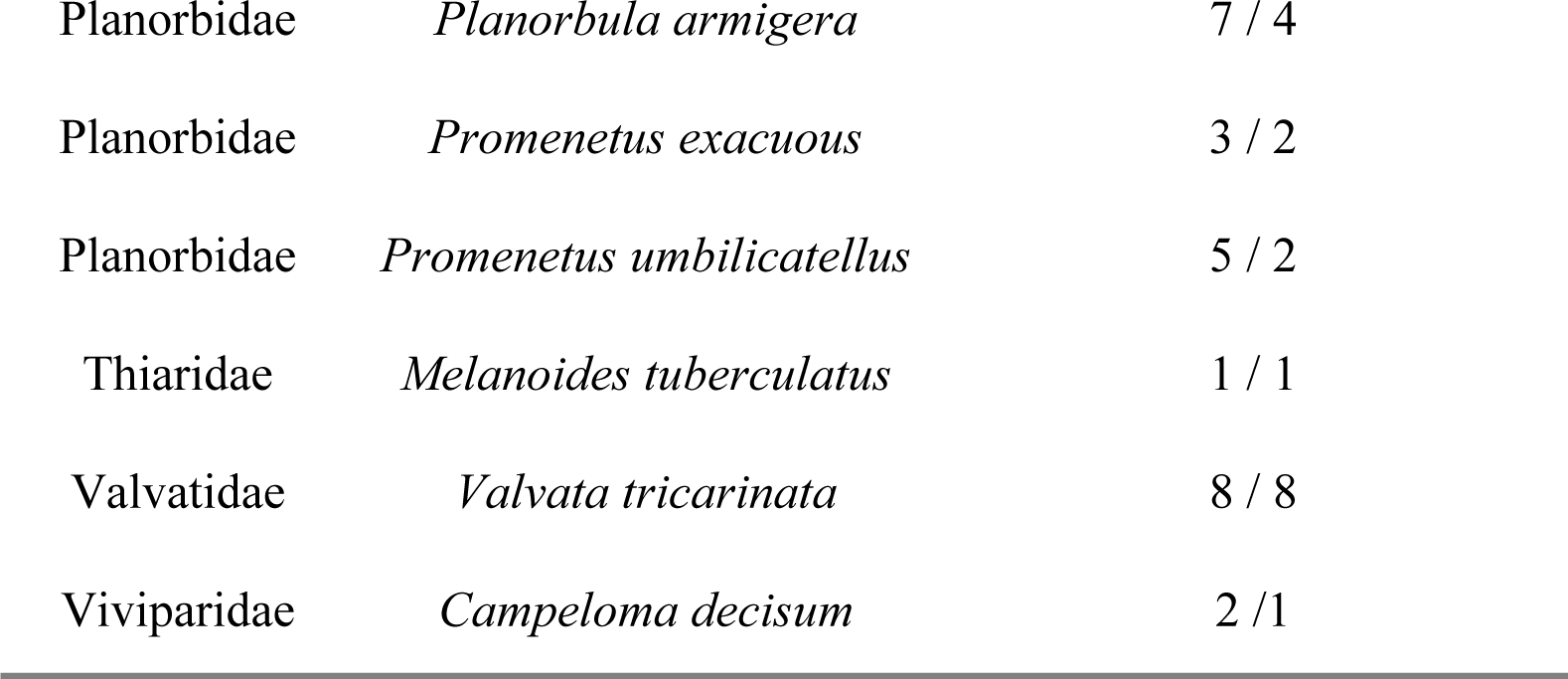
Records of freshwater gastropods in South Dakota based on literature and museum records.

| Family | Species | Literature & Museum<br>records:<br># Records / # Counties<br>(Comments in literature) |
| --- | --- | --- |
| Amnicolidae | *Amnicola limosus* | 3 / 3 |
| Hydrobiidae | *Cincinnatia integra* | 3 / 2 |
| Hydrobiidae | *Probythinella emarginata* | 5 / 5 |
| Lymnaeidae | *Lymnaea bulimoides* | 8 / 3 (over state) |
| Lymnaeidae | *Lymnaea caperata* | 17 / 8 (common over state) |
| Lymnaeidae | *Lymnaea catascopium* | 1 / 1 |
| Lymnaeidae | *Lymnaea elodes* | 16 / 10 (over State) |
| Physidae | *Aplexa elongata* | 1 / 1 |
| Physidae | *Physa acuta* | 10 / 5 (rare over state) |
| Physidae | *Physa gyrina* | 35 / 14 (abundant over state) |
| Physidae | *Physa jennessi* | 1 / 1 |
| Planorbidae | *Ferrissia rivularis* | 4 / 2 |
| Planorbidae | *Gyraulus crista* | 1 / 1 |
| Planorbidae | *Gyraulus deflectus* | 6 / 6 |
| Planorbidae | *Gyraulus parvus* | 11 / 8 |
| Planorbidae | *Helisoma anceps* | 6 / 2 (common over state) |
| Planorbidae | *Helisoma trivolvis* | 17 / 8 (Common over state) |
| Planorbidae | *Planorbula armigera* | 7 / 4 |
| Planorbidae | *Promenetus exacuus* | 3 / 2 |
| Planorbidae | *Promenetus umbilicatellus* | 5 / 2 |
| Thiaridae | *Melanoides tuberculatus* | 1 / 1 |
| Valvatidae | *Valvata tricarinata* | 8 / 8 |
| Viviparidae | *Campeloma decisum* | 2 / 1 |

**Table 2.** Conservation status based on species incidence of freshwater gastropods based on 98 recently samples sites throughout South Dakota. Historical records is the number of literature and museum records. Conservation status is based on quartile rank (see text).

| Species | Species Incidence | Conservation Status | Historical Records |
| --- | --- | --- | --- |
| *Physa gyrina* | 69 | S5 | 35 |
| *Lymnaea elodes* | 59 | S5 | 16 |
| *Gyraulus parvus* | 40 | S5 | 11 |
| *Helisoma trivolvis* | 39 | S5 | 17 |
| *Aplexa elongata* | 19 | S4 | 1 |
| *Promenetus exacuus* | 16 | S4 | 3 |
| *Lymnaea caperata* | 15 | S4 | 17 |
| *Valvata tricarinata* | 10 | S4 | 8 |
| *Helisoma anceps* | 6 | S3 | 6 |
| *Lymnaea stagnalis* | 6 | S3 | 1 |
| *Physa acuta* | 4 | S3 | 1 |
| *Lymnaea humilis* | 4 | S3 | 30 |
| *Physa jennessi* | 4 | S3 | 1 |
| *Ferrissia rivularis* | 2 | S2 | 4 |
| *Campeloma decisum* | 2 | S2 | 2 |
| *Amnicola limosus* | 2 | S2 | 3 |
| *Melanoides tuberculatus* | 2 | NA | 1 |
| *Planorbula armigera* | 1 | S1 | 7 |
| *Lymnaea bulimoides* | 0 | SH | 8 |
| *Gyraulus deflectus* | 0 | SH | 6 |
| *Probythinella emarginata* | 0 | SH | 5 |
| *Promenetus umbilicatellus* | 0 | SH | 5 |
| *Cincinnatia integra* | 0 | SH | 3 |
| *Gyraulus crista* | 0 | SH | 1 |
| *Lymnaea catascopium* | 0 | SU | 1 |

Based on the quartile method using incidence data I rank four species of indigenous snails as S2 or S1, *Ferrissia rivularis, Campeloma decisum, Amnicola limosus, Planorbula armigera* (Table 2). I exclude the non-indigenous *Melanoides tuberculatus* from ranking. I assign a conservation status of SH (possibly extirpated, historical listing only) or SU (unable to evaluate) to seven species listed historically but not observed in my recent surveys; *Lymnaea bulimoides, Gyraulus deflectus, Probythinella emarginata, Promenetus umbilicatellus, Cincinnatia integra, Gyraulus crista*, and *Stagnicola catascopium* (though see discussion on this species).

Distribution details for each species observed recently or historically is listed by family. Synonyms listed for each species are those used historically within South Dakota.

### Amnicolidae and Hydrobiidae

– The families Amnicolidae, Hydrobiidae and the Pomatiopsidae (the last not represented by any species herein) have overlapping descriptions and a history of misidentifications (Burch 1989). These families all contain tiny conical snails. The family Amnicolidae was often placed as a subfamily of Hydrobiidae, however, recent molecular work separates the families into separate clades (Wilke et al. 2001, Bouchet and Rocroi 2005). Lumping of historically listed species provides some clarity but many historic listings are still unclear. For example, some historic listings of *Amnicola cincinnatiensis* are actually *Probythinella emarginata* while others are *Cincinnatia integra* (Hershler 1996).

### *Amnicola limosus* (Mud Amnicola)

This small snail was listed in historical studies by Over (1915b) and Henderson (1927) with records from Brookings, Clay, and Codington counties. In recent sampling I observed this species from two sites, Jones Lake in Hand County and the Big Sioux River in Lincoln county. Regionally it appears common with records from all neighboring states examined (Cvancara 1983, Beetle 1989, Stewart 2006, Stephen 2015).

### *Amnicola walkeri* (Canadian Duskysnail)

This tiny snail was listed only by Over (1928) from Marshall county. No records of this species exist from adjacent states and it is out of range in South Dakota (Burch 1989). I did not observe this species in recent sampling and I treat the historic record as a misidentification.

### *Cincinnatia integra* (Midland Siltsnail)

Synonym: *Amnicola cincinnatiensis*. This small snail was listed in historical studies both by Over (1915b) and Henderson (1927) from Brookings, Moody, and Spink counties. The Museum of Comparative Zoology has records of this species from Turtle River in Spink County (MCZ 2014).

I did not observe this species in recent sampling. This species is recorded regionally from Nebraska (Stephen 2015), Iowa (Stewart 2006), and North Dakota (Cvancara 1983).

### *Probythinella emarginata* (Delta Hydrobe)

Synonyms: *Amnicola emarginata, Probythinella lacustris*. This tiny snail is listed by two historical studies from Codington, Clay, Moody, Roberts, and Brookings counties (Over 1915b, Henderson 1927). Burch (1989) placed it in South Dakota following Hibbard and Taylor (1960). I did not observe this species in recent sampling. The only regional listing is from Iowa (Stewart 2006). A recent systematic treatment recognizes a single species *Probythinella emarginata* among many synonyms (Hershler 1996).

Lymnaeidae.– There is wide systematic and taxonomic confusion among this family. Often placed in different genera according to size with the smallest placed into *Galba* or *Fossaria, Stagnicola* used for medium sized snails, and *Lymnaea* used for the largest (Burch 1989, Johnson *et al*. 2013). Dillon *et al*. (2013) suggests placing them all under the same genus *Lymnaea*. Confusion also exists among the many synonyms within this family, an area ripe for more molecular analysis. Most of the small Lymnaeids were synonimized by Hubendick (1951) under *Lymnaea humilis* including *L. parva* listed in South Dakota by Over (1915b). It seems likely that *Lymnaea elodes* is the conspecific elongate morph of *Lymnaea catascopium* as suggested by a gene tree of this family (Correa et al. 2010). Another species listed historically is *Lymnaea palustris*, which is restricted to Europe. I include *Lymnaea palustris* as a synonym of *L. elodes* following Burch (1989). One species listed historically by Over (1915b), *Lymnaea tyroni*, I could not resolve.

### *Lymnaea bulimoides* (Prairie Lymnaea)

Synonyms: *Lymnaea cokerelli, Lymnaea techella*. Over (1915b, 1928) list this species from Harding County and generally over the state. The Academy of Natural Sciences (ANS) collection includes records from Harding, Pennington, and Corson counties. The Museum of Comparative Zoology collections include a record of this species without a specific locality (MCZ). I did not observe this species in recent sampling. Regionally this species is listed from Nebraska and Wyoming (Hibbard and Taylor 1960, Beetle 1989). Given the taxonomic uncertainty, Stewart (2006) placed all historical listings of the small Lymnaeids (including *L. bulimoides)* together under *L. humilis*.

### *Lymnaea caperata* (Wrinkled Pondsnail)

Synonym: *Stagnicola caperata*. This species is listed by Over (1915b) and Henderson (1927) as well as Hibbard and Taylor (1960). It is considered common over the State (Over 1915b). Henderson (1927) lists three localities, Lake Campbell, Spring Creek, and Stony Butte in Brookings, Todd, and Lyman counties respectively. Hibbard and Taylor (1960) list localities from Perkins, Pennington and Lake counties. Museum collections include records from the additional counties of Clay, Corson, and Butte (ANP 2014). This species is sometimes considered under sampled due to its presence primarily in temporary wetlands (Jokinen 1992). In recent sampling I observed this species at 15 sites from 13 counties. Regional records are from Nebraska, North Dakota, and Iowa (Stephen 2015, Cvancara 1983, Stewart 2006).

### *Lymnaea catascopium* (Woodland Pondsnail)

Synonyms: *Galba apicina, Limnaea catascopium*. This snail is placed in South Dakota with a single record from Stanley County as *Galba apicina* (Baker 1911) following Binney (1865) who actually listed it as *Limnaea catascopium*. Baker thought this was a misidentification and altered it (Baker 1911). In addition to this problematic identification this species may be shown to be conspecific to *L. elodes* and several other medium size marsh snails (Correa *et al*. 2010). I did not observe this species in recent sampling. There are no regional listings of *Lymnaea apicina* but *Lymnaea catascopium* is recorded from Wyoming (Beetle 1989) and Iowa (Stewart 2006).

### *Lymnaea elodes* (Marsh Pondsnail)

Synonyms: *Stagnicola elodes, Lymnaea palustris*. Historical listings of this species are from two authors and two museum collections. Counties with records are Brookings, Clay, Corson, Deuel, Hamlin, Harding, Jones, Lyman, Moody, and Perkins (Over 1915, Henderson 1927, ANS, MCZ). Over (1915) also listed this species as “over the state”. In recent sampling I observed this species at 59 sample sites from 28 counties. There are regional records from each of the adjacent states of North Dakota, Wyoming, Iowa, and Nebraska (Cvancara 1983, Beetle 1989, Stewart 2006, Stephen 2015).

### *Lymnaea humilis* (Golden Lymnaea)

Synonyms: *Lymnaea obrussa, Lymnaea parva*. This snail is listed historically in the state by (Over 1915b) and the one museum collection (ANP). Records are from Butte, Corson, Clay, Deuel, Edmunds, Fall River, Harding, Lincoln, Pennington, and Perkins counties. Due to taxonomic uncertainly this nomen is used by some authors to encompass all tiny to small Lymnaeids (Stewart 2006). In recent sampling I observed this species at four sample sites from four counties, Beadle, Codington, Deuel, and Roberts. Regional listings of this species are from Nebraska, North Dakota, Wyoming, and Iowa (Stephen 2015, Cvancara, 1983, Beetle 1989, Stewart 2006).

### *Lymnaea stagnalis* (Great Pondsnail)

This species has consistent naming and no synonyms exist. This snail was listed only from Deuel county (Over 1915b). This species is considered common in northern North America from large water bodies (Burch 1989). In recent sampling I observed this species at six sample sites from Codington, Day, and Roberts counties. Listed regionally from a single location in Nebraska (Aughey 1877) it appears more common in North Dakota (Cvancara 1983) and northern Iowa (Stewart 2006).

Physidae. – I merge many nominal species of this family based on recent research using both traditional mating experiments (Dillon et al. 2002, Dillon and Wethington 2004) and molecular data (Wethington and Lydeard 2007). These works suggest that of 40 or so names of Physidae in North America there are perhaps eleven actual valid species. The two most abundant members of this family, *Physa gyrina* and *Physa acuta*, take on most of the nominal species. I follow the taxonomy suggested by Wethington and Lydeard (2007) and use a two-genus system, *Physa* and *Aplexa*, for this family. One species listed historically, *Physa humerosa*, not examined in recent systematic studies is placed under the subgenus *Costatella* by Burch (1989). All other species in this subgenus are merged with *Physa acuta* (Wethington and Lydeard 2007) therefore I consider this a synonym of *P. acuta* as well. *Physa warreniana* listed historically is not considered valid by Burch (1989) and has been treated as a subspecies of *P. sayi* (Walker 1906, Wu 2004-2005), which in turn is a synonym of *P. gyrina* (Wethington and Lydeard 2007). I could not resolve one species, *Physa crandilli*, (Over 1915b).

### *Aplexa elongata* (Lance Aplexa)

Synonym: *Aplexa hypnorum*. This species was listed by a single historical study from Deuel County (Over 1915b). In my recent sampling I observed this species at 19 sample sites from 12 counties. This species is common throughout the region with records from Nebraska, North Dakota, Wyoming, and Iowa (Stephen 2015, Cvancara 1983, Beetle 1989, Stewart 2006).

### *Physa acuta* (Bladder Physa)

Synonyms: *Physa ancillaria, Physa humerosa. Physa walkeri*. Over (1915b) recorded this snail from three counties Spink, Codington, and Lawrence, while Henderson (1927) recorded one site, Rapid Creek in Pennington County. The Museum of Comparative Zoology (MCZ) also houses specimens from Rapid Creek and the Academy of Natural Sciences of Drexel University (ANS) has records from Union and Washabaugh counties: Washabaugh County no longer exists having been split and merged into Jackson, Pennington, and Shannon counties. In my recent sampling I observed this species at four sample sites from four counties, Brookings, Codington, Fall River, and Kingsbury. This species appears common regionally being observed in North Dakota, Wyoming, Iowa, and Nebraska (Cvancara 1983, Beetle 1989, Stewart 2006, Stephen 2015).

### *Physa gyrina* (Tadpole Physa)

Synonyms: *Physa sayi, Physa warreniana, Physella gyrina*. This snail is listed as abundant over the entire state (Over 1915b). Specific records are from Deuel, Harding, Pennington, and Perkins counties (Over 1915, Henderson 1927). Museum collections contain records from the additional counties of Brookings, Clay, Codington, Custer, Fall River, Lawrence, Stanley, Union, Yankton, and the former county of Washabaugh (ANS, MCZ). This species has the highest incidence in my recent sampling being observed at 69 sample sites from 44 counties. This is a common species being listed in all regional states examined, North Dakota, Wyoming, Iowa, and Nebraska (Cvancara 1983, Beetle 1989, Stewart 2006, Stephen 2015).

### *Physa jennessi* (Obtuse Physa)

Synonym: *Physa skinneri*. There is a single museum record of this species from Deuel County (ANS). In my recent sampling I observed this species at four sample sites from four separate counties, Clark, Hamlin, Marshall, and Roberts. Regional observations are from Wyoming and Nebraska (Beetle 1989, Stephen 2015).

### Planorbidae

– This is the most specious family of freshwater snails (Strong et al. 2005). Some recent molecular systematic work has delved into a limited number of species in this family (Albrecht *et al*. 2007), but the most complete treatment is morphological (Hubendick and Rees 1955). I could not resolve two species of historically listed planispiral snails, *Planorbis vermicularis* and *Sementina crassilabris*. Both species are listed by Over (1915b). The fresh-water limpets, formerly in the family Ancylidae, are now placed within the Planorbidae (Bouchet and Rocroi 2005). Recent work has reduced the many possible historical species of fresh water limpets thought to inhabit North America to just three valid species throughout the entire U.S. (Walther et al. 2006, Walther and Burch 2010). One species listed historically, *Ferrissia tarda*, I could not resolve (Over 1928). The MCZ lists a specimen of genus *Ancylus* (now *Ferrissia*), from the Vermillion River, without other details (MCZ).

### *Ferrissia rivularis* (Creeping Ancylid)

Synonyms: *Ferrissia parallela*, *Ancylus parallelus*. This is a small freshwater limpet. Records of this species are from Deuel and Marshall counties (Over 1915b, 1928). In my recent sampling I observed this species at two sample sites from two separate counties, Pennington and Lawrence. This limpet is abundant in Iowa (Stewart 2006) and was observed at more than 40 sites in North Dakota (Cvancara 1983). It is also recorded from at least four counties in Wyoming (Beetle 1989).

### *Gyraulus crista* (Nautilus Ramshorn)

Synonym: *Segmentina cristyi*. This species is listed from a single “small pond” in Deuel county (Over 1915b). I did not observe this species in my recent sampling. Regionally records of this species are from the northern adjacent states of North Dakota and Wyoming plus a single record from Nebraska (Cvancara 1983, Beetle 1989, Taylor 1960). This species is abundant in the northern U.S. and Canada (Burch 1989).

### *Gyraulus deflectus* (Flexed Gyro)

Synonyms: *Planorbis deflectus, Planorbis hirsutus*. This species is listed by two studies (Over 1915b, 1928). Counties listed are Deuel, Clay, Perkins, and Marshall and the former counties of Washington and Washabaugh, which have been sliced up or merged into Jackson, Pennington and Shannon counties. I did not observe this species in my recent sampling. Regionally this species is recorded from Nebraska and Iowa (Aughey 1877, Stewart 2006).

### *Gyraulus parvus* (Ash Gyro)

Synonym: *Planorbis parvus*. This species is listed by Over (1915b) and one museum collection (ANS) from Brookings, Clap, Deuel, Hamlin, Harding, Lincoln, Moody, and Pennington counties. In my recent sampling I collected this species at 40 sample sites from 30 counties. Regionally it appears common with listings from all the neighboring states examined here, North Dakota, Wyoming, Iowa, and Nebraska (Cvancara 1983, Beetle 1989, Stewart 2006, Stephen 2015).

### *Helisoma anceps* (Two-ridged Ramshorn)

Synonym: *Planobula antrosa*. This species is listed in only one historical study as being observed over the state (Over 1915b). Museum records list this species from Perkins County and the former county of Washabaugh (ANS). In my recent sampling I collected this species at six sample sites from Gregory, Jones, Pennington, and Potter counties. Regionally it appears common with records from North Dakota, Wyoming, Iowa, and Nebraska (Cvancara 1983, Beetle 1989, Stewart 2006, Stephen 2015).

### *Helisoma trivolvis* (Marsh Ramshorn)

Synonym: *Planorbella trivolvis, Planorbis trivolvis*. Two studies list this species (Over 1915b, Henderson 1927). Records of this species also appear in museum collections (ANS, MCZ). Over (1915b) considered them common over the state. Specific counties with records are Brookings, Clay, Deuel, Hamlin, Jones, Lyman, Pennington, Perkins, Stanley, and the former county Wasabaugh (ANS, MCZ, Henderson 1927). In recent sampling I collected this species at 39 sample sites from 24 counties. Regional records exist from all adjacent states examined (Cvancara 1983, Beetle 1989, Stewart 2006, Stephen 2015).

### *Planorbula armigera* (Thick-lipped Ramshorn)

Synonym: *Sementina armigera*. This species is listed in South Dakota from two studies and one museum collection in Brookings, Clay, Deuel, and Moody counties (ANS, Over 1915b, Henderson 1927). I collected this species at a single sample sites from Hamlin county. This species is considered widely distributed in North America (Burch 1989). Regionally North Dakota, Iowa, and Nebraska have records of *P. armigera* (Cvancara 1983, Stewart 2006, Stephen 2015).

### *Promenetus exacuous* (Sharp Sprite)

Synonym: *Planorbis exacuus*. This snail is listed by a single study and one museum collection from Deuel and Edmunds counties (Over 1915b, ANS). In my recent sampling I collected this species at 16 sites from 10 counties.This species is common regionally being observed in all adjacent States examined (Cvancara 1983, Beetle 1989, Stewart 2006, Stephen 2015).

### *Promenetus umbilicatellus* (Umbilicate Sprite)

Synonym: *Planorbis umbilicatellus*. This snail is listed historically in South Dakota from Corson and Perkins Counties (ANS, Over 1915b) and is also included on a map (Hibbard and Taylor 1960, Fig 8 page 112). The map includes two sites in the northeast and one in the northwest, but specific locality information is not included. I did not observe this species in my recent sampling. Regionally it is listed from Iowa (Stewart 2006), North Dakota (Cvancara 1983), and Wyoming (Beetle 1989). The Iowa listing is historical and is from a single unspecified location (Stewart 2006). Burch (1989) lists *P. umbilicatellus* in Colorado but recent surveys failed to find it or the congeneric *P. exacuous* (Harrold and Guralnick 2010).

Thiaridae.– The Thiaridae are a tropical family common in the aquarium trade. Some members of this family, including *Melanoides tuberculatus*, are parthenogenic (Burch 1989).

### *Melanoides tuberculatus* (Red-rimmed Melania)

Synonym: *Melanoides tuberculata*. This species is not listed in historical studies and is not indigenous to South Dakota. This tropical species is known from just one area of South Dakota, the warm water streams around the town of Hot Springs in the southern Black Hills (Anderson 2004). In my recent sampling I collected this species at two sample sites both from Fall River County, one from Fall River itself and one from the adjacent Kidney Springs. There are records of this species from Colorado where it is similarly confined to warm springs (Harrold and Guralnick 2010).

Valvatidae.– Species within this family are considered common in northern North America (Clarke 1981, Jokinen 1992). The southern extent of their range is not well clarified but extends beyond South Dakota; *Valvata tricarinata* is found in Nebraska (Gugler 1969).

### *Valvata tricarinata* (Three-ridged Valvata)

Synonym: *Valvata winnebagoensis*. This species was observed in South Dakota by two researchers (Over 1915b, Henderson 1928). There are also records from two museums (ANS, MCZ). Specific records are from Brookings, Codington, Corson, Deuel, Hamlin, Marshall, Moody, and Roberts counties. I collected this species at 10 sample sites from 10 separate counties. Regionally it appears common with records from North Dakota (Cvancara 1983), Nebraska (Gugler 1969), and Iowa (Stewart 2006). This species is not listed in Wyoming but there are records of the congeneric *Valvata sincera* (Beetle 1989).

Viviparidae. – The Vivipariade are large ovoviviparous snails common in the northeast U.S. Historically several species within this family are recorded in adjacent Nebraska (Aughey 1877), however, only the range of *Campeloma decisum* is considered to be within this region (Burch 1989).

### *Campeloma decisum* (Pointed Campeloma)

Synonym: *Campeloma integrum*. Records of this species are from Clay County and the Vermillion River (Over 1915b). The species is most often observed in large lakes and slow-moving rivers and may be under sampled since it burrows in sand or mud (Jokinen 1983). In my recent sampling I collected this species at two sample sites from two separate counties, Lincoln and Union. In North America this species is considered common from South Dakota to the east coast (Burch 1989). Regional listings are from Iowa and Nebraska (Stewart 2006, Stephen 2015).

## Discussion

Historical records list 54 species names for freshwater gastropods within South Dakota. I reduce these nominal species to 25 valid species including one recent non-indigenous addition (Table 1). Reduction of the number of species from historical lists is primarily due to systematic revisions that lump species. Nineteen of the twenty-five species listed are pulmonates. This high pulmonate abundance is due to their ability to thrive in ephemeral wetlands throughout the state, which are particularly abundant in northeast of South Dakota.

Historically ten species appear relatively common and distributed broadly, more than five records in more than five counties, throughout South Dakota. Six of these species *Physa gyrina*, *Lymnaea elodes*, *Gyraulus parvus, Helisoma trivolvis, Lymnaea caperata*, and *Valvata tricarinata* have high incidence in the recent survey (Table 2). The other four species are *Lymnaea humilis, Lymnaea bulimoides, Physa acuta*, and *Gyraulus deflectus*. I observed a higher incidence of two species, *Aplexa elongata* and *Promenetus exacuous*, than expected based on historic records.

Species I tentatively list as S2 or S1, *Ferrissia rivularis, Campeloma decisum, Amnicola limosus, Planorbula armigera* (Table 2) should be of primary conservation concern. However, I suspect *Campeloma decisum* is more common than this list suggests due to its burrowing behavior making detection more difficult. The other species here are small which could also prevent detection, though other species in this size range (e.g. *Promenetus exacuous)* were readily observed.

### Problematic Species

*Lymnaea catascopium*, indeed the Lymnaiedae in general, is in a problematic taxonomic and systematic position. In South Dakota a single record accounts for both *Lymnaea catascopium* and *Lymnaea apicina. Lymnaea catascopium* might also be conspecific to *Lymnaea elodes* (Correa *et al*. 2010) and this is likely the case for *Lymnaea apicina* as well. Considering these three species I expect only one species is actual present in South Dakota and the single record of *L. catascopium/apicina* is a misidentification. If conspecificity is determined the name *L. catascopium* would take presidence. I choose not to treat them as a single species, or use the nomen *catascopium*, since the morphs in this region appear as *L. elodes* is described (Burch 1989). However, this family is known for a large amount of phenotypic plasticity (Brown 1985, Wullschleger and Jokela 2002, Brönmark *et al*. 2011) and is ripe for detailed molecular taxonomy.

*Physa jennessi* first listed in the region as *Physa skinneri*, which appear to be a synonym (Wethington and Lydeard 2007), was not listed in earlier studies under that name since it was not described until 1954 (Taylor 1954). *Physa jennessi*, described earlier, and thought to be restricted to Alaska and Canada (Dall 1919). This species is small and may be confused with young of other Physidae.

*Gyraulus parvus*, found to be common, is nearly identical morphologically to *Promenetus umbilicatellus* and *Gyraulus circumstriatus*. These species are thus candidates for identification errors and perhaps are also conspecific (an area ripe for molecular phylogenetic analysis). Two other species, *Cincinnatia integra* and *Probythinella emarginata*, have known historical misidentifications (Hershler 1996) and therefore I find it likely that only one and not both of these species either were or are present in the region.

*Gyraulus crista* appears rare within the region with records from just four sites within North Dakota (Cvancara 1983), two county records from Wyoming (Beetle 1989), and a single record from Nebraska (Taylor 1960). This species is more common to the north being abundant in central Canada (Prescott and Curteanu 2004).

### Records not found

I discovered no specific locality data for four species: *Ferrissia fragilis, Gyraulus circumstriatus, Pomatiopsis lapidaria*, and *Valvata sincera*, listed as being in South Dakota by a historical review (Johnson *et al*. 2013). Each of these species is listed regionally in at least some adjacent states and therefore the presence in South Dakota is probable. The limpets suffer from inadequate generic descriptions (Basch 1963), which led to historic confusion. This problematic area has recently been clarified with molecular data lumping most of the nominal species into just three species throughout the entire U.S. (Walther et al. 2006, Walther and Burch 2010). This modern clarification, however, doesn’t help clear up unresolved historic names within South Dakota. There is a single listing of the genus *Ancylus* from MCZ and the unresolved species *Ferrissia tarda* from Roberts County – each of which could be *Ferrissia fragilis*. Records for *F. fragilis* are from each of the adjacent states of Kansas, Iowa, Colorado, and Nebraska (Leonard 1959, Stewart 2006, Harrold and Guralnick 2010, Stephen 2015), and thus it is likely present in South Dakota. Records of *Pomatiopsis lapidaria* are from Kansas, Iowa, and Nebraska (Leonard 1950, Stewart 2006, Stephen 2015) thus it also appears to be a likely South Dakota inhabitant. This species, however, appears to require spring water habitats (Angelo *et al*. 2002) which limits its distribution. *Valvata sincera*, observed in the adjacent states of Wyoming, Colorado, and Nebraska (Beetle 1989, Harrold and Guralnick 2010, Stephen 2015) is another likely candidate for presence in South Dakota though it doesn’t appear abundant in any of the adjacent states. In fact, three of these possible inhabitants of South Dakota, *Ferrissia fragilis, Pomatiopsis lapidaria*, and *Valvata sincera* are considered rare and are listed as species of conservation concern in some states (Smith 1984, KSDWPT 2009).

The scarcity of both historical and recent records makes highly confident evaluation of species conservation status challenging. Additional surveys and monitoring are needed to better evaluate the geographic range, species diversity hotspots, critical habitat, and overall conservation status of freshwater snail species within South Dakota.

## Acknowledgements

I thank Patricia Freeman and Keith Geluso for edits to earlier versions of this manuscript. Thanks to the South Dakota Department of Game, Fish and Parks for providing funding over several seasons through their Wildlife Diversity Small Grants Program. Thanks to Rob Dillon and the Freshwater Gastropods of North America project for providing the impetus for this study.

